# Gradual relation between perceptual awareness, recognition and pupillary responses to social threat

**DOI:** 10.1101/2022.09.20.508721

**Authors:** Marta Poyo Solanas, Minye Zhan, Beatrice de Gelder

**Affiliations:** Department of Cognitive Neuroscience, Faculty of Psychology and Neuroscience, Maastricht University, Maastricht, The Netherlands; Department of Computer Science, University College London, London, UK

## Abstract

There is substantial evidence about affective stimulus processing outside awareness in healthy participants and brain-damaged patients. However, there are still methodological concerns mainly relating to the methods used to assess awareness. In two experiments, we investigated the processing of social threat in healthy participants by combining the continuous flash suppression paradigm and the perceptual awareness scale, a finer measure of perceptual awareness than dichotomous (seen/unseen) responses. Our behavioral results show a gradual relationship between emotional recognition and perceptual awareness. Recognition sensitivity was also higher for fearful than angry bodies for all visual awareness levels except for the perceptual unawareness condition where performance was at chance level. Interestingly, angry body expressions were suppressed for a shorter duration than neutral and fearful ones. Pupil dilation was a function of affective expression, the duration of suppression and the level of perceptual awareness. In conclusion, behavioral as well as pupillary responses showed a gradual relationship with perceptual awareness, and this relationship was influenced by the specific stimulus category.

## Introduction

Of all the external and internal information that is continuously being processed by the brain, some reaches consciousness, but most does not. Yet, information that does not reach the stage of subjective report, verbal or other, still influences behavior. This fact is widely acknowledged at least since the first findings on subliminal perception in the ’50s. However, the processes that ultimately determine which information does reach consciousness are not well understood. In affective neuroscience, one area of consensus is that stimuli with affective significance tend to get prioritized (Pessoa, 2010) in the sense of having a better chance to access conscious perception, possibly because of their behavioral relevance for the organism. On the other hand, many studies have shown that affective stimuli are being processed and influence behavior without having reached the stage of conscious perception. A functional explanation of non-conscious affect perception may be that affective stimuli are to some extent processed by possibly pre-wired adaptive behavioral circuits that do not necessitate conscious perception (LeDoux, 2012). One way or another, the role of consciousness in the perception of affective stimuli continues to be hotly debated, and evidence comes from different populations, including patients and healthy participants.

Patients with cortical blindness following lesions in their primary visual areas sometimes preserve the ability to discriminate visual stimuli presented to their blind field even when being unaware of the stimulus (Weiskrantz, 1990; Weiskrantz, Warrington, Sanders, & Marshall, 1974). This phenomenon is called “blindsight” (Weiskrantz, 1990) and has been studied extensively in the field of visual perception and consciousness, initially for low level visual stimulus attributes such as the direction of motion (Ter Braak, Schenk, & Van Vliet, 1971). Later studies turned to stimuli with high behavioral relevance like facial and body expressions and showed that these images can also be processed outside conscious awareness, as found with behavioral (de Gelder, Vroomen, Pourtois, & Weiskrantz, 1999) as well as neuroimaging methods (Burra, Hervais-Adelman, Celeghin, de Gelder, & Pegna, 2019; de Gelder & Hadjikhani, 2006; Morris, de Gelder, Weiskrantz, & Dolan, 2001). Physiological measures have also been used to establish affective blindsight. For example, increased pupil dilation has been observed for fearful facial expressions as compared to happy expressions for seen but also for unseen conditions in blindsight patients (Tamietto et al., 2009). However, the visual system of these patients may have developed compensatory strategies; hence, it remains an open question whether similar phenomena can be observed in the intact brain.

In healthy participants, several methods have been used to render (emotional) stimuli invisible, such as the continuous flash suppression (CFS) paradigm (Tsuchiya & Koch, 2005). In CFS, the target stimulus is made invisible by presenting it with low-contrast to one eye, while the other eye is presented with a dynamic noise made of colorful patterns. The resulting interocular competition causes the conscious percept of the target stimulus to be suppressed by that of the colorful mask. This method has been increasingly used because, in comparison to previous approaches such as masking, it creates a stronger suppression and a more stable non-conscious perception (Yang, Brascamp, Kang, & Blake, 2014). Another advantage of this method is that it comes with different variants, such as priming (quantifies the effect of the invisible prime on the visible target) or breaking from suppression (quantifies the suppression time of the experimental stimulus) (for reviews see Stein, Hebart, & Sterzer, 2011; Yang et al., 2014).

Previous research in healthy participants, some of it using CFS, has indeed shown that emotional expressions have a special status among other visual stimuli, especially those signaling threat. For example, fearful faces break from suppression faster than other emotional expressions (Gray, Adams, Hedger, Newton, & Garner, 2013; Stein, Seymour, Hebart, & Sterzer, 2014; Tsuchiya, Moradi, Felsen, Yamazaki, & Adolphs, 2009; Yang, Zald, & Blake, 2007), although this difference did not reach significance in some studies (Sterzer, Hilgenfeldt, Freudenberg, Bermpohl, & Adli, 2011; Zhan, Hortensius, & de Gelder, 2015). Angry facial expressions, on the contrary, break from suppression slower than fearful and neutral faces (Gray et al., 2013; Zhan et al., 2015). For body expressions, shorter suppression times have been found for angry compared to both fearful and neutral body postures, with fearful body postures being the last category to break from suppression (Zhan et al., 2015). These findings suggest that fearful and angry bodies may be processed differently, despite both expressions signaling threat. A possible explanation of these findings may be that angry body expressions convey a direct threat signal while fearful body expressions are ambiguous about the cause of fear (Pichon, de Gelder, & Grèzes, 2009). This pattern is particularly interesting when considering the different behavioral responses that these two threatening expressions trigger with, on one hand, anger initiating flight and/or responses and, on the other hand, fear triggering freezing behavior (Mello et al., 2022; Roelofs, 2017).

Although considerable evidence has been gathered in the last decades about non-conscious processing in both blindsight patients and healthy participants, perceptual processing without accompanying awareness has long been controversial. This has to do with ongoing discussions about consciousness or awareness itself as well as with methodological debates about criteria to assess it. For example, the vast majority of studies explicitly measuring subjective perceptual awareness have used a dichotomous measure (i.e., yes-no responses), which may be inadequate for capturing weak conscious experiences and the level of participant’s awareness of the stimulus (Mazzi, Bagattini, & Savazzi, 2016; Overgaard, Fehl, Mouridsen, Bergholt, & Cleeremans, 2008). Recently, more fine-grained measures of perceptual awareness have been developed because they presumably better capture intermediate levels of perceptual awareness. One of these is the perceptual awareness scale (PAS), developed by Ramsoy and Overgaard (2004) (Ramsøy & Overgaard, 2004). This scale aims to reflect different states of subjective perceptual awareness of a stimulus by having four response alternatives: (1) “no experience”, (2) “brief glimpse”, (3) “almost clear experience” and (4) “clear experience”. The use of this scale provides the opportunity to address the findings from blindsight patients and healthy subjects, by differentiating genuine forms of blindsight from degraded conscious vision (Mazzi et al., 2016). In this regard, several studies using PAS to measure stimulus visibility have reported chance performance in objective forced-choice discrimination tasks during perceptual unawareness (Hesselmann, Darcy, Rothkirch, & Sterzer, 2018; Lähteenmäki, Hyönä, Koivisto, & Nummenmaa, 2015; Lamy, Alon, Carmel, & Shalev, 2015; Lamy, Carmel, & Peremen, 2017; Lohse & Overgaard, 2019; Peremen & Lamy, 2014; Ramsøy & Overgaard, 2004; Tagliabue, Mazzi, Bagattini, & Savazzi, 2016).

Another issue in consciousness research is that it is often difficult to separate the measurement methods from the (implicit) theory about consciousness (e.g., Sandberg, Timmermans, Overgaard, & Cleeremans, 2010; Wierzchoń, Asanowicz, Paulewicz, & Cleeremans, 2012; Wierzchoń, Paulewicz, Asanowicz, Timmermans, & Cleeremans, 2014). A dichotomous measurement with forced-choice methods fits the notion that awareness is a matter of all or nothing. But the use of fine-grained measures fits the notion that perceptual awareness is a graded rather than an all-or-none phenomenon. Extensive efforts have gone into trying to resolve this debate, resulting in considerable evidence supporting both accounts. Therefore, it is still unclear whether perceptual awareness is a gradual or a dichotomous phenomenon and how this relates to affective stimuli.

In this study, we investigated the processing of emotional body expressions by using two CFS paradigms in combination with the PAS. In the first experiment, body postures expressing anger, fear or a non-emotional expression (i.e., neutral) were randomly presented to the left or right visual field of participants’ non-dominant eye, while a colorful dynamic noise mask was shown to the dominant eye. Participants first performed a two-alternative forced-choice task (angry/fear vs. neutral) and subsequently rated their visual experience of the stimulus according to the perceptual awareness scale (Ramsøy & Overgaard, 2004). In a second experiment, we measured the breaking from suppression time of these same bodily expressions and the subjective visual experience (i.e., PAS rating) at the moment of breaking from suppression. For both experiments, changes in pupil size of participants’ non-dominant eyes were recorded using an eye-tracking device. This experimental design allowed us to answer four main questions. First, it allowed us to investigate how perceptual awareness relates to objective stimulus presence, and more specifically, to objective emotional recognition performance and pupillometry. Secondly, it also provided the chance to investigate whether this relation is gradual or dichotomous. Thirdly, it gave us the opportunity to assess affective processing in conditions of perceptual unawareness. Finally, it allowed the investigation of whether behavioral and pupillometric measures are sensitive to the different stimulus conditions. Particularly, whether there are differences in the processing of direct and indirect threatening body expressions.

## Materials and Methods

### Participants

Seventy-six healthy participants were recruited for two experiments. Their data were included in the analyses if (1) they showed successful and stable merging of the stimuli (i.e., saw one rectangular frame instead of two; see *Stimuli, task design and experimental procedure* section), (2) performed the task correctly and (3) their visual perception through the non-dominant eye was not completely suppressed by that of the dominant eye (i.e., body stimuli occasionally escaped suppression). The latter criterion was adopted to allow the investigation of body expression processing outside conscious awareness but also at different levels of perceptual awareness. Only the data of 30 participants satisfied these criteria and were used for the analysis of experiment 1 (mean age = 21.8 years; age range = 18-27 years; 19 females; 3 left-handed participants, all of them female). Of these 30 participants, only the data of 11 participants were used for experiment 2 due to issues in recording behavioral responses (N = 11; mean age = 20.7 years; age range = 18-24 years; 7 females; one left-handed participant, which was female). All participants had normal or corrected-to-normal visual acuity, normal stereo and color vision and no medical nor any psychiatric or neurologic disorders. All participants were naïve to the CFS paradigm and remained unaware of the aim and the experimental set-up of the study. Participants received credit points or monetary reward after their participation. The study was performed with the understanding and written consent of each participant, in accordance with the Declaration of Helsinki, and all procedures followed the regulations of the Ethical Committee at Maastricht University.

### Stimuli, task design and experimental procedure

The experiment consisted of two sessions performed on separate days. In the first session, participants performed an eye dominance test (6 min), followed by two practice runs of experiment 1 (4 min each) and six runs of experiment 1 (10-12 min each). The second session consisted of the six runs of experiment 1 and two runs of experiment 2 (16 min each). The emotional body expressions for experiment 1 differed from one session to the other. One of the sessions used angry and neutral conditions (AN-session) while the other fearful and neutral ones (FN-session). The condition order for the sessions of experiment 1 was randomized across participants. Experiment 2 was always performed in the second session. The total duration of the study was approximately five hours.

The tasks were presented in MATLAB vR2007a (MathWorks, Natick, MA, USA) using Psychtoolbox 3.0.11 (Brainard & Vision, 1997; Pelli & Vision, 1997) on an LCD screen (Iiyama prolite b2483hsu, resolution = 1920 x 1080 pixels, screen width = 53cm, screen height = 30cm, refresh rate = 60Hz) under constant and controlled dim-light conditions. Participants rested their heads on a chinrest placed in front of the screen at a distance of 99 cm. A cardboard panel was situated between the chinrest and the screen, dividing the screen into two halves and ensuring that each eye would not receive information from the contralateral side of the screen. The dichoptic presentation was achieved using a pair of prism glasses, which projected the ipsilateral image to each eye’s field of view center by bending the light from the screen. To facilitate the fusion of the images perceived by each eye, two black rectangular frames with a fixation cross in the center were placed next to each other (404 pixels apart, 6.45° visual angle) on a grey background (RGB value = 128, 128, 128). The specific diopter of the prism glasses (diopter = 6) was chosen based on the visual angle between the two rectangular frames (Schurger, 2009). This experimental setup ensured that each of the participant’s eyes only perceived one of the rectangular frames at the center of the screen (Schurger, 2009). Therefore, the right side to the fixation cross within the rectangle corresponds to the right visual field of the participant while the left side to the left visual field, for both eyes. Apart from the instructions specific to each part of the study, participants were asked to keep their heads as still as possible throughout the experiments, remain fixated on the fixation cross, and not to blink within the CFS period of each trial if possible. In addition, each part of the experiment would start only after the participant reported a stable perception of a single rectangular frame.

### Eye dominance test

For the eye-dominance test, ten identities of neutral faces (half female) taken from the Radboud Face Database (Langner et al., 2010) were used. These stimuli (318×212 pixels, 5.08°x3.39° visual angle) were presented to one eye in the center of the rectangular frame (318×212 pixels, 5.08°x3.39° visual angle, 10 pixels of frame thickness) while the dynamic mask pattern (318×212 pixels, 5.08°x3.39° visual angle) flashing at 10Hz was shown to the other eye. Each stimulus was randomly presented three times to each eye, giving a total of 60 trials. Each trial had a duration of 3s, consisting of 1s of a gradual ramping up of the stimulus contrast from 0% to full contrast, which was next maintained for 1s, and then diminished to 0% contrast over 0.5s, followed by a 0.5s blank period. Subsequently, participants reported whether they saw or did not see a face by pressing one out of two keys (“J” for seen, “K” for unseen) with the right hand. The dominant eye was then defined as the eye that perceived the highest amount of seen trials. In the cases where the amount of seen trials was equal between both eyes, eye dominance was assigned randomly. This was the case for one participant included in the analysis.

### Experiment 1

In experiment 1, participants’ dominant eye was presented with a flickering colorful mask (318x352 pixels, 5.08°x5.62° visual angle) covering the entire rectangular frame (318x352 pixels, 5.08°x5.62° visual angle, 10 pixels of frame thickness; see **Figure 1A**). The colorful mask consisted of 600 unique patterns flashing randomly at 10Hz, each composed of small overlapping rectangles. Participants’ non-dominant eyes were presented with a static body posture (318x182 pixels, 5.08°x2.91° visual angle) to the left or right side of a center fixation cross in a randomized order, while no stimulus was presented to the other side. The body stimuli were developed and validated in the lab (see Stienen & de Gelder, 2011). The stimulus set consisted of angry, fearful and neutral (i.e., opening door) body expressions that had the information of the face removed. Opening door was chosen as the neutral condition to control for action and implied motion information present in the emotional conditions. Eight actor identities (half females) were used across the three conditions and were presented in greyscale on a grey background (RGB value = 128, 128, 128).

**Figure 1.**
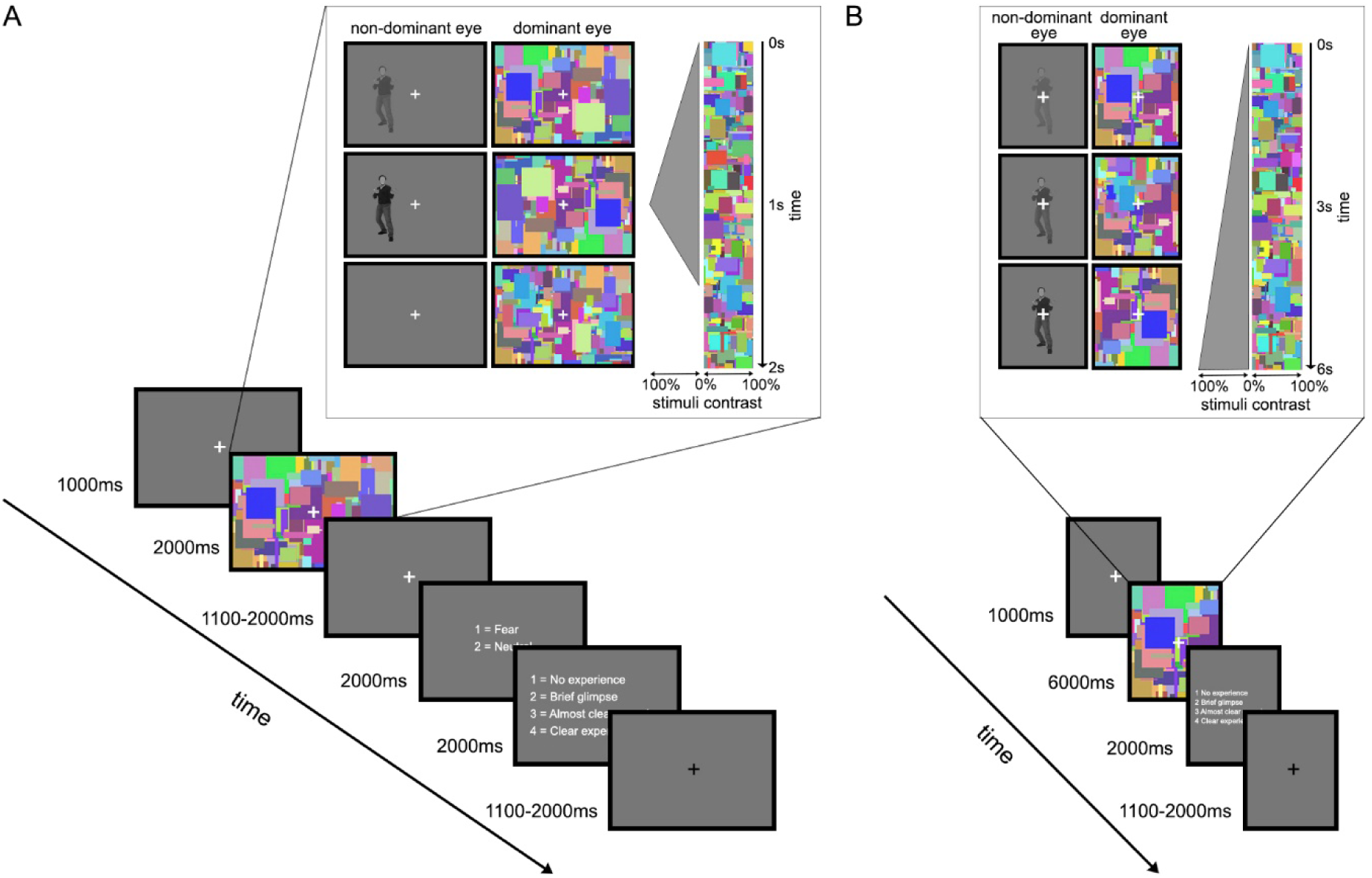
Schematic view of a trial presentation sequence in both experiment 1 and 2. **A)** Experiment 1. After a 1s-fixation, the 2s-CFS trial started with 1s of a gradual ramping up of the body stimulus contrast from 0% to full contrast, followed by the contrast diminishment to 0% within 0.5s and 0.5s blank period (see content within frame). After a jittered fixation period, participants were required to make two active responses, each within a 2s window: a two-alternative forced-choice task (angry/fear vs. neutral) and the rating of their visual experience of the stimulus according to the Perceptual Awareness Scale (PAS). The inter-trial-interval was jittered (1.1-2s, 100ms steps) and the average trial duration was 10s. **B)** Experiment 2. Each trial consisted of a 1s-fixation period, followed by a CFS presentation consisting of a gradual ramping up of the body stimulus contrast from 0% to full contrast (see content within frame). Participants were required to press “J” as soon as they perceived something in the noise, which terminated the CFS presentation that otherwise lasted 6s. Subsequently, participants were required to rate their visual experience of the stimulus according to the PAS. The inter-trial-interval was jittered (1.1-2s, 100ms steps).

The start of each trial was indicated by the change in color of the fixation cross from black to white and it remained white throughout the trial. In each trial, a 1s-baseline fixation period was followed by a 2s-CFS presentation, starting with 1s of a gradual ramping up of the stimulus contrast from 0% to full contrast, followed by the contrast diminishment to 0% within 0.5s and a 0.5s blank period. The gradual increase of the stimulus contrast was performed to decrease the likelihood of the target stimulus escaping suppression. Subsequently, participants were required to make two active responses. The first response was a two-alternative forced-choice task (2AFC; angry vs. neutral or fearful vs. neutral depending on the session) by pressing one out of two keys (“J”, “K”). Next, participants rated their visual experience of the stimulus according to the perceptual awareness scale (Ramsøy & Overgaard, 2004) by pressing one out of four keys: “no experience” (PAS1, key “J”), “brief glimpse” (PAS2, key “K”), “almost clear experience” (PAS3, key “L”) and “clear experience” (PAS4, key “;”). Participants understood this scale as a question about the clarity of the percept or about different degrees of visibility, not about the quality of the awareness as a separate attribute. To facilitate the responses, the possible answers were shown on the screen during the response window, with numerical values to the left indicating the finger/key and the corresponding descriptions to the right. The key assignment was randomized on a trial basis for the emotional categorization task but remained constant for the visual experience task due to the higher number of response options. Both responses were required, even when participants failed to see anything in the noise. In those cases, participants were asked to guess the emotional body posture and rate their visual experience as “no experience”. Both responses were always performed with the right hand within a 2s-response window, respectively. The inter-trial-interval was jittered (1.1-2s, 100ms steps) and the average trial duration was 10s. Each run consisted of 64 trials, 32 per condition (four repetitions for each of the eight stimulus identities). Therefore, a total of 384 trials were obtained in each of the two sessions (six runs each), giving a final amount of 192 trials per participant for the angry category, 192 for the fearful category and 192 + 192 for the neutral condition. One participant only participated in the AN session (i.e., 6 runs of the FN session are missing). One participant completed the six runs of the AN session but only performed two runs of the FN session. Two participants completed the six runs of the FN session but only completed 3 and 5 runs, respectively, of the AN session.

In order to gain familiarity with the PAS ratings, participants performed two practice runs before the actual experiment 1. These practice runs used facial (Radboud Face Database; Langner et al., 2010) and body expressions (Stienen & de Gelder, 2011) depicting happiness, fear, anger and a non-emotional expression (opening door in the case of the body postures and a neutral expression for faces). Three male actor identities were used that differed from the ones shown in the main experiment. Each stimulus was presented twice, once per run, giving a total of 48 trials. Each trial followed a procedure as in the main experiment, with the exception that participants only needed to rate their visual experience with the PAS without emotionally categorizing the stimulus.

### Experiment 2

Experiment 2 used a CFS paradigm that measured the breaking from suppression time of the same body postures used in Experiment 1. A change in the fixation color from black to white indicated the start of each trial one second before the beginning of the CFS period and remained white throughout the trial **(Figure 1B)**. In contrast to Experiment 1, the fixation period was followed by a CFS presentation where the body stimulus was presented centrally, and its contrast was gradually ramped up from 0% to full contrast without ramping down. Participants were required to press “J” as soon as they perceived something in the noise, whether that was a “brief glimpse”, the full stimulus or anything in between. Pressing “J” terminated the CFS presentation that otherwise lasted 6s. After the CFS part, the PAS scale appeared in the screen for 2s, indicating participants to rate their visual experience of the stimulus at the moment of pressing “J” in the same manner as in Experiment 1. If participants did not perceive anything in the noise, they were instructed to report “no experience” (i.e., PAS1). Responses were always performed with the right hand. The inter-trial-interval was jittered (1.1-2s, 100ms steps). Each run consisted of 120 trials, 40 per condition (five repetitions of each of the eight stimulus identities). Therefore, a total of 240 trials were obtained (two runs), giving a final amount of 80 trials for each category (i.e., angry, fearful and neutral). No practice runs were performed for this experiment.

### Recording of pupil size data

The eye movements and pupil diameter of participants’ non-dominant eye were recorded during Experiment 1 and 2 using a monocular pupil-tracking infrared camera operated by ViewPoint EyeTracker® Software (version 2.9.2,5; Arrington Research Inc., Scottsdale, AZ, USA) with a sampling rate of 90.5 Hz. Gaze position was computed using ViewPoint’s non-linear algorithm based on the dark pupil and pupil-glint vector methods. Pupil diameter was calculated using ViewPoint’s Ellipse method. A calibration test was performed before each run. In this test, participants were required to fixate at the center of twelve targets represented as green squares on the stimulus monitor, appearing one at a time on an imaginary 12-point grid in the half side of the screen that corresponded to the non-dominant eye. The measured eye position signals for each of these target squares were then used to map optimally the location of the gaze into the subject’s GazeSpace coordinates. Before calibration, the pupil and corneal reflection were isolated with appropriate threshold settings. To facilitate the fixation during the calibration test, the dominant eye was covered.

### Analysis of behavioral data

#### Experiment 1

To understand possible perceptual differences between stimulus categories, Signal Detection Theory measures (SDT) (Green & Swets, 1966; Tanner & Swets, 1954) were used across the visual awareness ratings. In SDT, when having a two-alternative forced-choice task, participants’ performance is described by four parameters: hits (H), misses (M), correct rejections (CR) and false alarms (FA). In the current 2AFC task, hits refer to “anger” (AN session) or “fear” (FN session) responses on signal trials (i.e., trials where one of these target stimulus categories was displayed) whereas misses refer to an incorrect “neutral” response on signal trials. Correct rejections indicate a correct “neutral” response on noise trials (i.e., when the non-target stimulus, or neutral, was displayed) and false alarms refer to noise trials where incorrect “anger/fear” responses were given.

According to SDT, participants’ decisions depend on both the perceptual sensitivity (d’) to discriminate between stimulus categories and the response criterion or bias (c), which is the tendency to favor one stimulus type over another independently of sensitivity. Sensitivity is defined as the distance between the target and the noise distribution means in standard deviation units and it is usually computed by subtracting the z-transformed hit and false alarm rates (see **Equation 3**). To account for ceiling effects, Snodgrass & Corwin (1988) proposed the use a modified form of hit (H’) and false alarm rates (FA’) (**Equation 1 & 2**) (Snodgrass & Corwin, 1988). A sensitivity value of zero indicates inability to differentiate between the emotional category (i.e., fear or anger) from the non-emotional one (i.e., neutral condition). Higher values are indicative of a better sensitivity in making the distinction. Criterion bias refers to the distance, in standard deviation units, between the response criterion and the neutral point where responses are neither favored towards “emotional stimuli” nor “neutral stimuli”. It is calculated by summing the z-transformed hit and false alarm rates and then multiplying the result by 0.5 (**Equation 4**) (Macmillan, 1993; Snodgrass & Corwin, 1988; Stanislaw & Todorov, 1999; Tamietto, Geminiani, Genero, & de Gelder, 2007). Positive criterion bias values indicate a conservative response criterion (i.e., improbability of reporting the presence of the emotional stimulus regardless of its actual presence) whereas negative values show a liberal criterion (i.e., bias toward reporting the presence of an emotional stimulus).

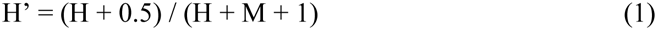

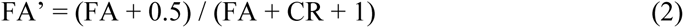

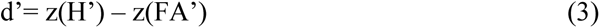

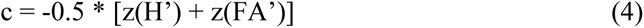

D’ and c scores were calculated for each of the PAS ratings and sessions, respectively. Before their calculation, outliers were removed based on reaction times (RTs) that deviated more than 3.5 times the standard deviation from the mean within a subject. Trials without a response for one or both ratings as well as those in which gaze shifts deviated more than 2° from the fixation cross were also excluded from the analysis. A total of 746 out of 11136 trials (6.7%) were excluded from the FN session and a total of 749 out of 10624 trials (7.1%) were excluded from the AN session. These excluding criterions were applied to all the analyses performed for Experiment 1. Subsequently, d’ and c values were analyzed in SPSS (version 22.0), respectively, using a linear mixed model procedure. Both models included as within-subject factors Emotion (two levels: Fear, Anger) and Visual Awareness (four levels: PAS1, PAS2, PAS3 and PAS4) and used the Toeplitz covariance matrix for repeated measures based on Akaike information criterion (AIC) values (Akaike, 1974). The weighted least squares method was used to account for violations of homoscedasticity. The Sidak method was employed in all analyses of the current study to correct for post-hoc comparisons. In addition, one sample t-tests against zero were performed to compare the d’ and c scores to chance level, separately for each session and visual awareness level.

To further investigate whether perceptual sensitivity presents a gradual or an “all-or-none” relationship to perceptual awareness, two linear mixed models were fitted to the participant data. These two models differed in the type of predictors used and were performed separately for the anger and fearful conditions within each subject. In the gradual model, the predictors modelled a linear relationship between sensitivity values and the PAS ratings whereas in the “all-or-none” (dichotomous) model, the PAS1 predictor was set to 0 while the rest of the PAS levels were set to 1, describing an “all-or-none” relationship between perceptual sensitivity and perceptual awareness. The final model selection was performed following the model with the significantly lowest Bayesian information criterion (BIC) value (Stone, 1979), which indicates better fit. To do so, the BIC values resulting from each model fitting were entered into a linear mixed model analysis with within-subjects factor Model (two levels: gradual and dichotomous) and Emotion (two levels: anger and fear) and the Toeplitz covariance matrix. Following a significant effect of Model, but not a significant interaction, a paired t-test was performed between the coefficient estimates of the anger and fearful models to assess how different the model slopes and intercepts were between emotions.

Finally, the reaction times of the emotional categorization task and the visual awareness ratings were analyzed using a linear mixed model procedure. The analysis of the RTs of the emotional categorization task included as within-subject factors Session (two levels: FN, AN) and SDT measures (four levels: H, M, FA, CR) and used the Unstructured covariance matrix for repeated measures. Two outliers were removed from this analysis (single data-points within the whole sample) based on their standardized residuals resulting in a model with a significantly better fit. The analysis of RTs of the visual awareness task included as within-subject factors Session (two levels: FN, AN) and Visual Awareness (four levels: PAS1, PAS2, PAS3 and PAS4). Three observations (single data-points within the whole sample) were removed from this analysis based on their standardized residuals resulting in a model that now met the assumptions of normality and homoscedasticity. This analysis used the Toeplitz covariance matrix for repeated measures.

#### Experiment 2

For experiment 2, trials in which the RTs of the visual awareness task deviated more than 3.5 times the standard deviation from the mean within a subject were excluded from further analyses. Trials without visual awareness rating and trials in which participants “broke from suppression” but reported not seeing the body stimulus (i.e., PAS1) were also excluded. In addition, trials with gaze shifts deviating more than 2° from the fixation cross were excluded from the analysis. These criteria resulted in the rejection of a total of 78 out of 2640 trials (2.95%). Subsequently, breaking from suppression times were analyzed using a linear mixed model procedure with within-subject factors Emotion (three levels: Neutral, Fear, Anger) and Visual Awareness (three levels: PAS2, PAS3 and PAS4). Apart from the above-mentioned criteria, PAS1 was not included as a Visual Awareness level since no breaking from suppression occurred when participants reported “no experience” of the stimulus. This analysis used the Toeplitz covariance matrix for repeated measures and the weighted least squares method to account for violations of homoscedasticity. One observation (single data-point within the whole sample) was removed from this analysis based on their standardized residuals resulting in a model with significantly better fit.

Reaction times of the visual awareness task were also analyzed using a linear mixed model procedure with within-subject factors Emotion (three levels: Neutral, Fear, Anger) and Visual Awareness (four levels: PAS1, PAS2, PAS3 and PAS4) and the Toeplitz covariance structure. Two outliers were removed from this analysis (single data-points within the whole sample) based on their standardized residuals resulting in a model with a significantly better fit.

### Pre-processing and analysis of pupil size data

#### Pre-processing of pupil size data

Pupil size data were inspected for various artifacts with custom code in MATLAB (version R2020a; The MathWorks Inc., Natick, MA, USA). Pupil size samples that were outside a biologically feasible range were rejected (e.g., pupil size smaller than 2mm in diameter). Samples that presented a large variation in absolute pupil size with respect to adjacent samples (“speed outliers”) were removed (Kret & Sjak-Shie, 2019). In addition, samples within 50ms adjacent to gaps in the data were removed to avoid artifacts resulting from blinks (e.g., pupil size misestimation due to eyelid occlusion). Gaps were defined as contiguous sections of missing data larger than 75ms. Remaining speed outliers and outliers that were four standard deviations from the mean were rejected. After these steps, missing data were interpolated linearly. The resulting data were smoothed with a zero-phase 10th-order low-pass filter with cut-off frequency at 4Hz (Jackson & Sirois, 2009).

Trials that required linear interpolation to more than 50% of the data corresponding to the baseline (0-1s), CFS (1-3s) and fixation (3-4.5s) periods, respectively, were excluded from further analysis. This criterion did not apply to the response period (4.5-8.5) since these data were not used for further analysis and participants were allowed to blink during this time. Apart from the linear interpolation constraint, the trial exclusion criteria applied to the behavioral analyses of Experiment 1 and 2, respectively, were also applied for the pupillometry analyses (see *Analysis of behavioral data* section). After applying these criteria, a total of 1232 out of 11136 trials (11%) were excluded from the FN session and a total of 1299 out of 10624 trials (12%) were excluded from the AN session of Experiment 1. In Experiment 2, 221 trials out of 2640 were excluded (8%). Finally, each data sample of the remaining trials was normalized by the average pupil size recorded during 500ms of the baseline fixation period preceding each CFS period, respectively.

#### Analysis of pupil size data

##### Experiment 1

To investigate changes in pupil dilation over time, the baseline-corrected pupil size data were analyzed by dividing the 2s-CFS period in consecutive 500ms time bins. Therefore, average estimates of four 500ms time bins were obtained for each trial. The last 500ms of the 2s-CFS period (Time bin 4) was included in the analysis despite no body expression being presented during that time bin (see **Figure 1A**), so that any potential late effects in pupil dynamics could be investigated. Subsequently, pupil size data were analyzed with a Generalized Estimating Equations (GEE) model due to severe violations of normality. This model included the within-subject factors Emotion (three levels: Neutral, Fear and Anger), Visual Awareness (four levels: PAS1, PAS2, PAS3 and PAS4) and Time bin (four levels: 0-500ms, 500-1000ms, 1000-1500ms, 1500-2000ms). The Unstructured covariance matrix was used for repeated measures based on Quasi Information Criterion (QIC) values (Pan, 2001). The weighted least squares method was used to account for violations of homoscedasticity.

Furthermore, we also investigated whether pupil dilation presented a gradual or an “all-or-none” relationship to perceptual awareness with a similar procedure to the one employed for the sensitivity values (see *Experiment 1* subsection within the *Analysis of behavioral data* section). The only difference was that this analysis used an Unstructured covariance matrix. The pupil size data used for this analysis consisted of the average pupil diameter of the 2s-CFS period after baseline correction.

##### Experiment 2

The average estimates of the baseline-corrected pupil diameter data were calculated for the 200ms before the breaking from suppression point. This was performed separately for each visual awareness rating and emotion. Subsequently, pupil size data were analyzed with a linear mixed model including the within-subject factors Emotion (three levels: Neutral, Fear and Anger) and Visual Awareness (three levels: PAS1, PAS2 and PAS3) and Breaking from Suppression Time as a covariate (in its centered form). The compound symmetry covariance matrix was used for repeated measures. Two observations (single data-points within the whole sample) were removed from this analysis based on their standardized residuals resulting in a model with better fit.

## Results

### Behavioral results

#### Experiment 1

To understand behavioral differences in the processing of anger and fearful body expressions, a linear mixed model procedure was performed separately on two signal detection theory measures: sensitivity and criterion bias (see *Analysis of behavioral data* section). The analysis on sensitivity showed significant main effects of Emotion (F(1, 59.79) = 18.56, p < .001) and Visual Awareness (F (3,86.50) = 92.52, p < .001) as well as a significant Emotion*Visual Awareness interaction (F(3,86.90) = 7.57, p < .001), indicating a significant increase in sensitivity as a function of visual experience with the exception of PAS1 and PAS2 for anger (see **Figure 2A**). Participants also displayed a significantly higher sensitivity for fear than for anger in all visual experience levels except when participants reported not seeing the body stimulus. In addition, sensitivity values differed from the chance level (value of 0) for both angry and fearful body expressions in all Visual awareness ratings, with the exception of PAS1 for both emotions (**Figure 2A**).

**Figure 2.**
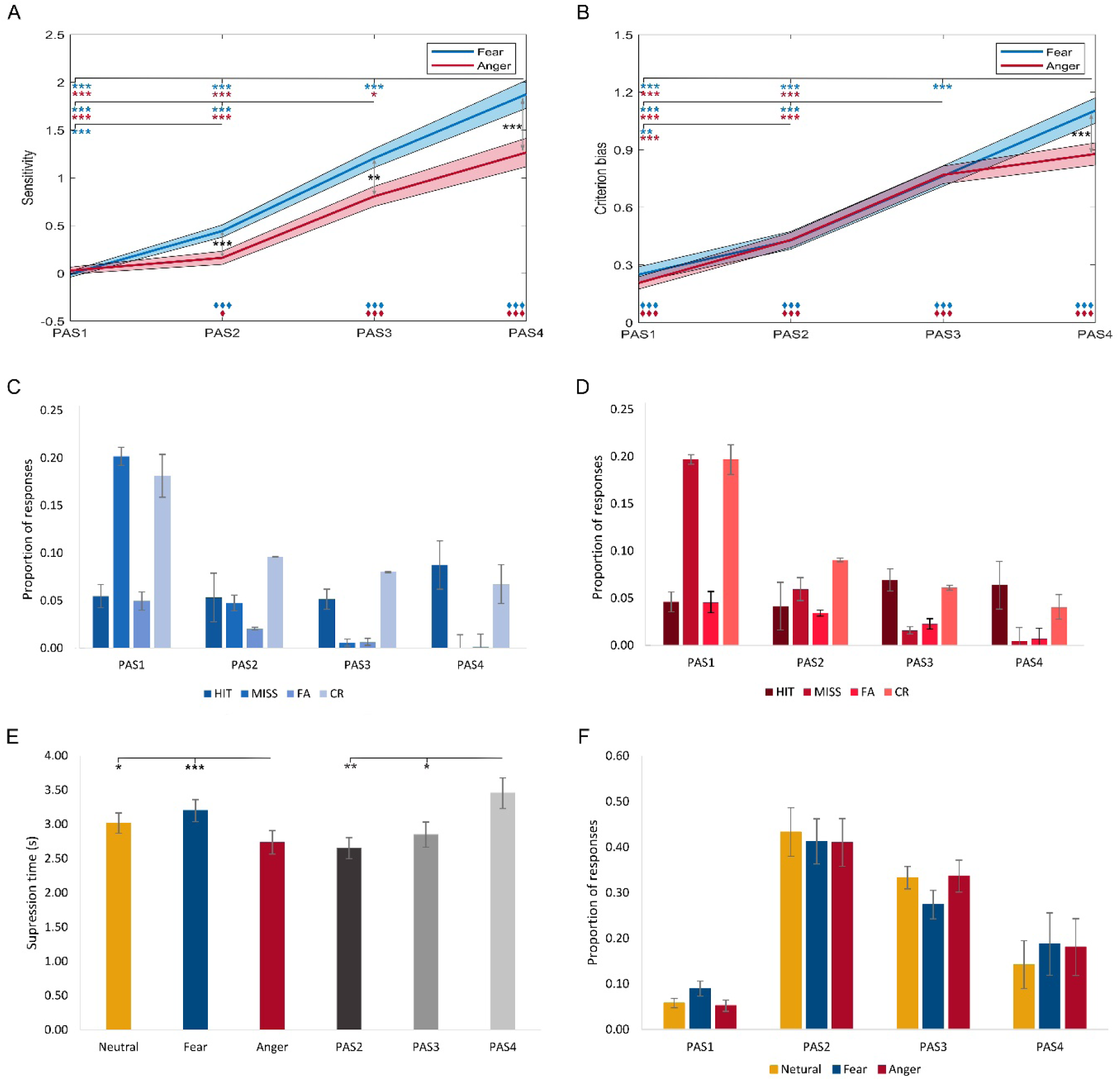
Overview of behavioral results of experiment 1 and 2. **A)** Average sensitivity values across PAS levels for both the FN and AN sessions of experiment 1. The reported values are estimated marginal means; **B)** Average criterion bias values across PAS levels for both the FN and AN sessions of experiment 1. The reported values are estimated marginal means; **C)** Average proportion of responses separated by PAS and SDT measures (i.e., Hit, Miss, FA, CR) for the FN session of experiment 1; **D)** Average proportion of responses separated by PAS and SDT measures for the AN session of experiment 1; **E)** Average suppression times (in seconds) for each body expression and visual awareness rating of experiment 2. The reported values are estimated marginal means; **F)** Average proportion of responses separated by the type of visual experience and emotion of experiment 2. Error bars and shadowed areas indicate standard error from the mean. Asterisks in red denote significant differences across PAS levels for anger while blue asterisks for fear (A, B). Black asterisks denote significant differences between fear and anger (A, B, E) or between PAS levels (E). Rhombi in red denote significant difference from zero for anger while blue rhombi for fear (A, B). */τ: p < .05; **/ττ: p < .01; ***/τττ: p < .001. ***Abbreviations***: AN: anger/neutral; Correct rejection (CR): neutral stimuli categorized as neutral; False alarms (FA): neutral stimuli categorized as emotional; FN: fear/neutral; Hit: emotional stimuli correctly categorized; Miss: emotional stimuli categorized as neutral; PAS: perceptual awareness scale: PAS1: “no experience”, PAS2: “brief glimpse”; PAS3: “almost clear experience”, PAS4: “clear experience”.

To better understand the relationship between perceptual sensitivity and perceptual awareness, two models were fit into the data that modelled either a linear or an “all-or-none” relationship. The final model selection was performed following the model with the significantly lowest BIC value, indicative of better fit (see **Table S1** in the Supplementary Information). This analysis yielded a significant main effect of Model (F(1,19.04) = 73.91, p < .001), showing a significantly better fit for the linear model (M = 0.17) than the dichotomous one (M = 5.52). There was no significant main effect of Emotion (F(1,28.22) = 1.40, p = .246) nor a significant Model*Emotion interaction (F(1,35.81) = 0.87, p = .359). Further analyses revealed that the model for the fearful body conditions had a significantly bigger slope (M = 1.76, SE = 0.17) than the model for angry body conditions ((M = 1.12, SE = 0.16; t(28) = -2.29, p = .030), indicating a bigger increase in sensitivity in response to increases of perceptual awareness for the fearful body expressions. No differences were found between emotional conditions regarding model intercepts (t(28) = -.16, p = .877).

The analysis of criterion bias values yielded a significant main effect of Emotion (F(1,43.00) = 4.73, p = .035) and Visual Awareness (F(3,55.83) = 73.64, p < .001) and a significant Emotion*Visual Awareness interaction (F(3,81.62) = 3.81, p = .013). Participants displayed a significantly more conservative response criterion bias (i.e., higher criterion bias values) as the visual experience of the stimulus became clearer (see **Figure 2B**). There was, however, no significant difference in the response bias between “almost clear” (i.e., PAS3) and clearly seeing angry bodies (i.e., PAS4). In addition, a significantly more conservative response criterion was found for fear (M = 1.10, SE = 0.07) than for anger (M = 0.88, SE = 0.06) during “clear experience” of the stimulus, but not at any other visual awareness level. Criterion bias values differed from zero at all visual awareness levels for both emotions, including PAS1 (p < .001; **Figure 2B**).

The analysis of the RTs of the emotional categorization task showed a significant main effect of SDT (F(3,28.64) = 4.75, p = .008), indicating faster responses when correctly categorizing emotional body expressions (H; M = 0.85; SE = 0.03) than when neutral body expressions were correctly categorized (CR; M = 0.88; SE = 0.03) or erroneously categorized as emotional (FA; M = 0.91; SE = 0.03) and also marginally faster than when incorrectly reporting emotional expressions as neutral (M; M = 0.90; SE = 0.03). No significant main effect was found for Session (F(1,27.80) = 0.34, p = .566) nor for the Session*SDT interaction (F(3,27.86) = 0.46, p = .714) (see **Table S2** in the Supplementary Information for estimated marginal means and standard errors).

The linear mixed model procedure on the RTs of the visual awareness ratings yielded a significant main effect of Visual Awareness (F(3,33.66) = 7.27, p = .001), indicating faster responses when participants reported not seeing anything (PAS1; M = 0.58, SE = 0.03) than when reporting seeing a “brief glimpse” (PAS2; M = 0.65, SE = 0.03), an almost clear perception of the stimulus (PAS3; M = 0.71, SE = 0.03) or a clear perception of the stimulus (PAS4; M = 0.66, SE = 0.03). No significant main effect was found for Session (F(1,37.60) = 0.17, p = .682) nor for the Session*Visual Awareness interaction (F(3,40.70) = 0.33, p = .801) (see **Table S2**).

#### Experiment 2

The analysis of breaking from suppression times showed significant effects of Emotion F(2,21.96) = 11.91; p < .001) and Visual Awareness (F(2, 16.94) = 7.31; p = .005). Breaking from suppression times were faster for angry (M = 2.74; SE = 0.17) than neutral (M = 3.02; SE = 0.15) and fearful (M = 3.20; SE = 0.16) body expressions (**Figure 2E**). In addition, longer suppression times were observed when participants reported having a “clear experience” of the stimulus (PAS4; M = 3.45; SE = 0.22) in comparison to when they had an “almost clear” experience (PAS3; M = 2.85; SE = 0.18) or just saw a “brief glimpse” (PAS2; M = 2.65; SE = 0.15) at the moment of breaking from suppression (**Figure 2E**). The interaction between Emotion and Visual awareness was non-significant F(4,31.67) = 1.11; p = .368) (see **Table S3** in the Supplementary Information for estimated marginal means and standard errors).

The analysis on the reaction times of the visual awareness ratings revealed no significant main effects for Emotion (F(2,10.74) = 2.81; p = .104) or Visual awareness (F(3,21.10) = 0.83; p = .493), nor a significant interaction between the two (F(6,25.91) = 0.43; p = .855).

### Pupillary results

#### Experiment 1

A generalized estimating equation model was performed to investigate pupil size differences between fearful, angry and neutral body expression perception under different visual awareness conditions over time. This analysis showed a significant main effect of Time bin (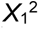 = 138.44, p < .001). There was also a marginally significant main effect of Visual awareness (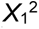 = 7.44, p = .059), showing that pupil size at PAS3 (M = -0.14, SE = 0.03) was significantly bigger than at PAS2 (M = -0.19, SE = 0.03) and marginally bigger than at PAS1 (M = -0.21, SE = 0.04) (**Figure 3**). In addition, there was a significant Visual awareness*Time bin interaction (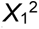 = 20.82, p = .013) and Emotion*Visual awareness*Time bin interaction (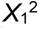 = 75.41, p < .001), showing that when not being aware of the stimulus (i.e., PAS1), pupil size was larger in Time bin 1 than in Time bin 2 & 3 for all emotions (see **Table S4** in Supplementary Information for estimated marginal means and standard errors). Pupil size at PAS1 was also bigger in Time bin 1 than in Time bin 4 for neutral bodies only and it was bigger in Time bin 4 than in Time bin 3 for neutral and angry bodies. For “brief glimpse” (i.e., PAS2), pupil size was larger in Time bin 1 than in Time bin 2 & 3 for all emotions. Pupil size at PAS2 was also bigger in Time bin 4 than in Time bin 3 for neutral and fearful bodies. When having an almost clear perception of the stimuli (i.e., PAS3), pupil size was larger in Time bin 1 than in Time bin 2 for all emotions, and also than in Time bin 3 for neutral and fearful bodies. In addition, pupil size at PAS3 was also bigger in Time bin 4 than in Time bin 3 for all emotions. Finally, when having a clear experience of the stimulus (i.e., PAS4), pupil size was larger in Time bin 1 than in Time bin 2 for neutral and fearful bodies and also than in Time bin 3 for neutral bodies only. Within each Time bin, no differences within or between emotions were found between PAS levels. There was no significant main effect of Emotion (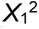 = 0.48, p = .785) as well as no significant Emotion*Visual Awareness interaction (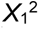 = 3.21, p = .782) nor Emotion*Time bin interaction (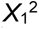 = 0.46, p = .998).

**Figure 3.**
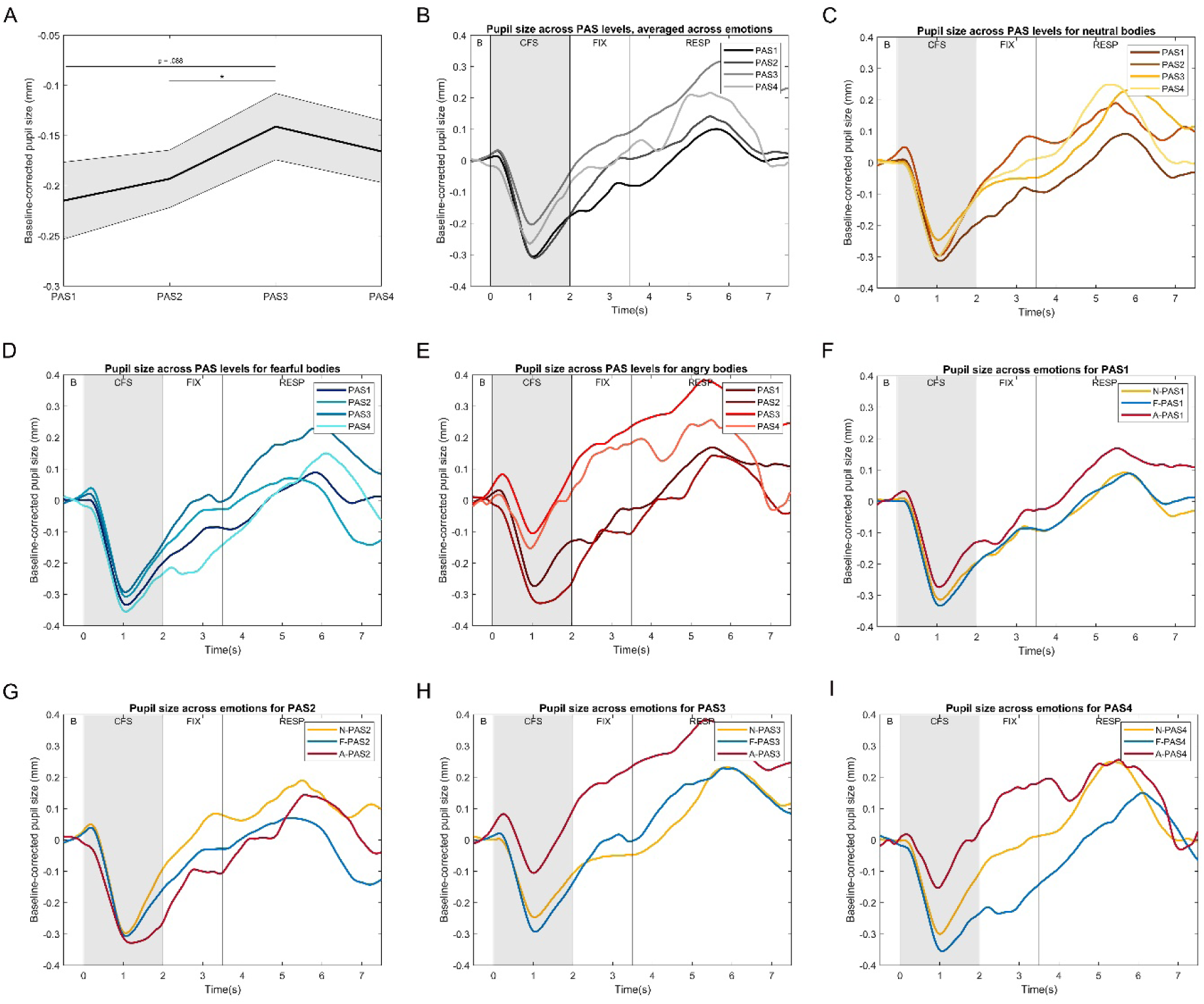
Pupil size changes over time across PAS levels and emotions of experiment 1. **A)** Average pupil size across PAS levels. The reported values are estimated marginal means. Asterisk denotes significant differences across PAS levels (*: p < .050). Shadowed area indicate standard error from the mean; **B)** Pupil size across PAS levels, averaged across emotions; **C)** Pupil size across PAS levels for neutral bodies; **D)** Pupil size across PAS levels for fearful bodies; **E)** Pupil size across PAS levels for angry bodies; **F)** Pupil size across emotions for PAS1; **G)** Pupil size across emotions for PAS2; **H)** Pupil size across emotions for PAS3; **I)** Pupil size across emotions for PAS4. ***Note***: pupil size time courses in B-I have been smoothed (movmean = 60) for visualization purposes. ***Abbreviations***: A: anger; B: baseline period; CFS: CFS period; F: fear; N: neutral; RESP: response period; PAS: perceptual awareness scale: PAS1: “no experience”, PAS2: “brief glimpse”; PAS3: “almost clear experience”, PAS4: “clear experience”.

Similar to the analysis conducted with the sensitivity values, the relationship between pupil diameter and perceptual awareness was investigated by fitting the pupillary data into a linear or an “all-or-none” model. There was no significant main effect of Model (F(1,25.88) = 1.05, p = .316) nor a significant Model*Emotion interaction (F(1,26.55) = .182, p = .835). The main effect of Emotion was marginally significant (F(1,22.35) = 3.08, p = .066). Following the positive trend observed between pupil size and the first three levels of perceptual awareness, a similar analysis was performed with only these three levels of perceptual awareness. This analysis yielded a significant main effect of Model (F(1,35.90) = 5.01, p = .032), showing that the gradual model (M = -11.97, SE = 1.04) fitted the data better than the dichotomous one (M = -10.80, SE = 1.04). A significant main effect of Emotion (F(2,52.24) = 3.88, p = .027) was found, revealing a significantly better fit for neutral bodies (M = -13.48, SE = 1.27) than angry ones (M = -9.76, SE = 1.32). The Model*Emotion interaction was not significant (F(2,50) = 0.06, p = .943).

#### Experiment 2

The comparison of the average estimates between emotions and visual awareness ratings as a function of the breaking from suppression time (BST; the covariate) showed a significant main effect of Visual Awareness (F(2,65.91 = 3.19, p = .048), indicating a significantly larger pupil size for PAS4 (M = -0.09, SE = 0.06) than PAS3 reports (M = -0.22, SE = 0.06) (**Figure 4A**). Also, there was a marginally significant main effect of Emotion (F(2,65.96 = 2.82, p = .067) and a marginally significant Emotion*Breaking from Suppression Time interaction (F(2,64.80 = 2.46, p = .093), indicating that the effect of the body expression on pupil size depends on the breaking from suppression time. To further investigate this, differences in pupil size between emotions were analyzed at three levels of the covariate: low, mean and high (low = early BST; mean = average BST; high = late BST). Mean corresponds to the average centered version of the original covariate which was calculated by subtracting the covariate mean from each individual score, and thus is zero. The low and high covariate levels were determined by adding or subtracting one standard deviation (calculated from the original covariate) to the cantered mean (Preacher, Curran, & Bauer, 2006; Preacher & Hayes, 2004). The results of these analyses revealed a marginally bigger pupil size for neutral (early BST: M = -0.09, SE = 0.06; average BST: M = -0.10, SE = 0.06) than fearful body expressions (early BST: M = -0.20, SE = 0.06; average BST: M = -0.20, SE = 0.06) only for early (p = .080) and average (p = .100) breaking from suppression times (**Figure 4B**). There was no significant main effect of Breaking from suppression time (F(1,74.00) = 0.03, p = .875), Emotion*Visual Awareness interaction (F(4,64.88) = 0.65, p = .633), Visual Awareness interaction*Breaking from suppression time interaction (F(2,68.58) = 0.12, p = .884) nor Emotion*Visual Awareness interaction*Breaking from suppression time interaction (F(4,65.34) = 1.82, p = .135).

**Figure 4.**
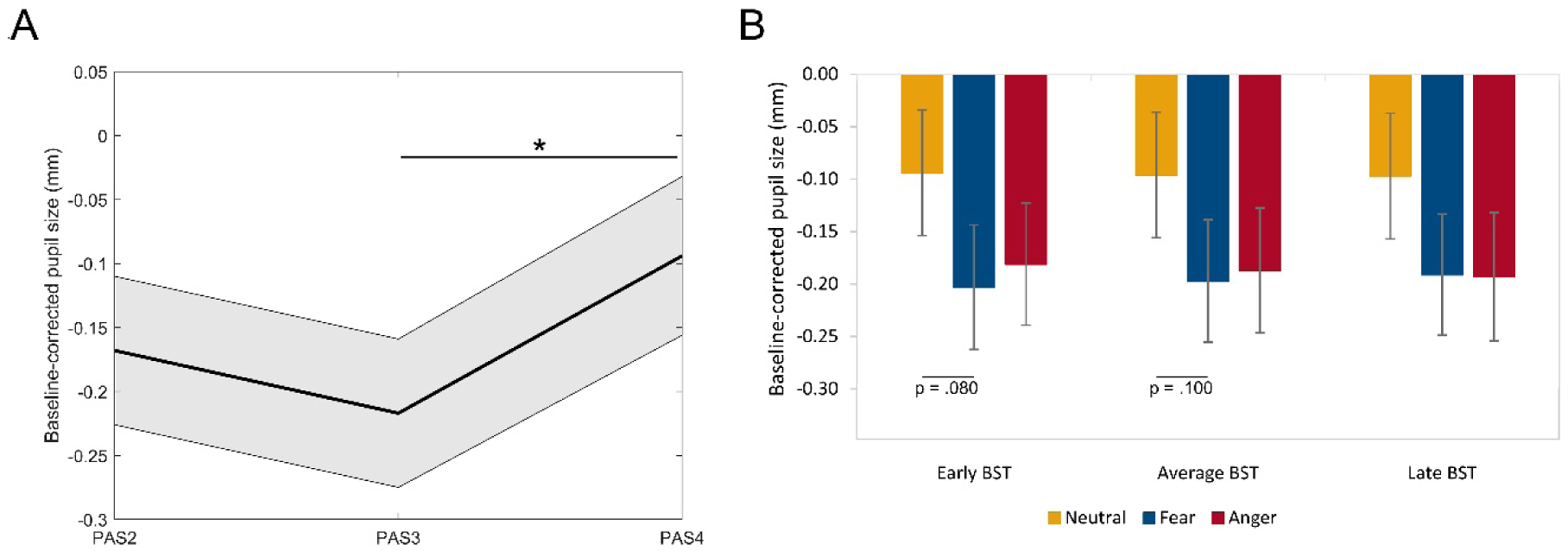
Results from the pupil size analyses of Experiment 2. **A)** Average pupil size across PAS levels. The reported values are estimated marginal means. Shadowed area indicate standard error from the mean. Asterisk denotes significant differences across PAS levels (*: p < .050); **B)** Average pupil size separately for different breaking from suppression times and emotions. The reported values are estimated marginal means. Error bars indicate standard error from the mean. ***Abbreviations***: BST: breaking from suppression time. PAS: perceptual awareness scale: PAS2: “brief glimpse”; PAS3: “almost clear experience”, PAS4: “clear experience”.

## Discussion

The aim of the current study was to investigate the relationship between perceptual awareness, emotion recognition and pupillary responses using a finer measure of perceptual awareness rather than a dichotomous task. We showed a gradual relationship between perceptual awareness and recognition sensitivity and no evidence of perceptual discrimination during perceptual unawareness. In addition, we observed a gradual relationship between pupillary responses and perceptual awareness, which was modulated by emotion and the time of breaking from suppression. Finally, we observed that anger and fearful body expressions were processed differently despite both expressions signaling threat.

### Perceptual awareness is gradual

The behavioral results of both experiment 1 and 2 support the graded account of perceptual awareness as participants rated their visual experiences using all levels of the measurement scale (see **Figure 2C** and **Figure 2D**) and sensitivity ratings were statistically better described by a gradual model than a dichotomous one. Furthermore, visual awareness correlated to emotional recognition performance, as higher emotional recognition sensitivity was observed with a clearer visual experience of the body stimulus (**Figure 2A**). This finding is in agreement with previous masking studies investigating the relation of different levels of visual awareness to the perception of simple features such as color or shape (Lähteenmäki et al., 2015; Overgaard, Rote, Mouridsen, & Ramsøy, 2006; Overgaard & Sandberg, 2012; Ramsøy & Overgaard, 2004; Sandberg & Overgaard, 2015; Sandberg et al., 2010; Wierzchoń et al., 2014; Windey, Vermeiren, Atas, & Cleeremans, 2014). Most importantly, it also supports recent findings showing a dependency between the degree of visual awareness and higher-level object and semantic perception, including facial expressions (Lohse & Overgaard, 2019) as well as other different classes of emotional stimuli (Lähteenmäki et al., 2015). Our study extends these findings by reporting a graded perceptual awareness account for body expressions, and even outside foveal vision, using a continuous flash suppression paradigm.

With regards to pupil size, no clear evidence for a gradual or a dichotomous account was found when taking into account all PAS levels. Although the analyses of Experiment 1 pointed to an overall positive trend between perceptual awareness and pupil size (**Figure 3A**), only significant pupil dilation differences were found between “almost clear” experience (i.e., PAS3) and both “brief glimpse” (i.e., PAS2) and perceptual unawareness (i.e., PAS1). In addition, pupil size seemed to be smaller when clearly seeing the body stimulus (i.e., PAS4) in comparison to an almost clear perception (i.e., PAS3), which could be the reason for the lack of gradual model preference. In fact, when model preference was evaluated considering only the first three levels of PAS, pupil dilation was then significantly better described by a gradual model than a dichotomous one. These results can be explained when taking into account the different functions attributed to pupillary reflexes. For example, pupil constriction has been related to increased visual acuity, important when we want to distinguish fine details (Mathôt, 2018). Conversely, pupil size dilations have been reported in situations when optimal vision is of essence, such as in the presence of faint stimuli (Mathôt, 2018). A larger pupil results in a bigger amount of light entering the eye, and with that, a greater amount of visual information. Thus, here, pupil dilation may have helped achieve higher visual sensitivity when subjective stimulus visibility was poorer, whereas pupil constriction may have facilitated finer evaluation of the body stimulus when its perception was clear (PAS4). Yet, the amount of pupil dilation in subjectively ambiguous conditions (i.e., PAS1, PAS2, PAS3) may still depend on subjective awareness, with bigger dilations as subjective awareness increases. It is important to note here that the changes in pupil dilation occurred in response to subjective awareness and not physical stimulus presence, as body stimuli were always presented with the same contrast pattern across trials (**Figure 1A**).

Taken together, the current results indicate that pupillary responses may serve different functions in response to different levels of subjective perceptual awareness, independently of the physical stimulus visibility. These results seem in disagreement with studies suggesting that pupillary responses are an objective measure of low- and high-level visual processing independent of conscious awareness (for reviews see Binda & Murray, 2015; Mathôt & Van der Stigchel, 2015). However, there is increasing evidence of cognitive influences on pupil responses (Bárány & Halldén, 1948; Brenner, Charles, & Flynn, 1969; Fahle, Stemmler, & Spang, 2011; Kimura, Abe, & Goryo, 2014; Lowe & Ogle, 1966; Naber, Frässle, & Einhäuser, 2011). Of particular interest here is the study by Naber and colleagues (2011), reporting that pupil dilation reflects perception rather than the physical stimulus, but also that binocular rivalry is a gradual phenomenon (Naber et al., 2011).

On a related issue, breaking from suppression at later time points was related to a clear perception of the body stimulus (i.e., PAS4) while earlier breaking from suppression times was associated with a limited body perception (i.e., PAS2 and PAS3) (**Figure 2E**). This could be explained by the fact that the contrast of the body stimulus increased gradually over time in Experiment 2, where breaking from suppression times were investigated (**Figure 1B**). Perceptual awareness at the time of breaking from suppression was also linked to pupil dilation, with larger pupil dilations for clear (i.e., PAS4) than almost clear (i.e., PAS3) experiences (**Figure 4A**). This contrasts with the pupillary findings reported in experiment 1, where pupil size was bigger for PAS3 than PAS4 (**Figure 3A**). A possible explanation may be that in Experiment 2, pupil size changes were only investigated at the moment of breaking from suppression while PAS reporting in Experiment 1 was not confined to that moment but to the whole CFS period. Therefore, it may be that pupil dilation has a different time course before, after and at the moment in which the body stimulus enters conscious awareness.

### Non-conscious processing of body expressions

Previous studies in both blindsight patients and healthy participants have provided evidence of emotion perception outside conscious awareness. However, in the current study no behavioral evidence of emotional processing outside perceptual awareness was observed, as recognition sensitivity for both angry and fearful body expressions did not differ from chance performance when participants reported not having seen the body stimulus (see **Figure 2A**). A possible reason of these divergent results could be that the majority of studies reporting non-conscious emotional processing used facial expressions (Tamietto & de Gelder, 2010), which may have different processing and detection mechanisms than bodies. For example, Zhan and colleagues (2015) found different suppression time patterns depending on whether the emotion was conveyed by a face or a body stimulus (Zhan et al., 2015). An important issue is also the low-level properties of the stimuli. For example, previous CFS experiments have reported that the faster breaking from suppression of fearful faces was directly related to the contrast of the eye region (Gray et al., 2013; Yang et al., 2007).

Another explanation for this discrepancy resides in the fact that previous studies have mostly used dichotomous measures (“yes/no”, “seen/unseen” answers), disregarding possible intermediate levels of perceptual awareness. It could be that the modulation observed during the “unseen” condition in those studies is a mixture between genuine forms of blindsight and degraded conscious vision (i.e., “no experience” and “brief glimpse”, respectively, in PAS) (Mazzi et al., 2016). Indeed, several studies measuring stimulus visibility with PAS have reported chance performance on objective forced-choice discrimination tasks during perceptual unawareness (e.g., Hesselmann et al., 2018; Lamy et al., 2015; Lamy et al., 2017; Peremen & Lamy, 2014; Ramsøy & Overgaard, 2004; Tagliabue et al., 2016) including studies investigating facial expression processing (Lähteenmäki et al., 2015; Lohse & Overgaard, 2019).

Consistent with the behavioral data, no clear evidence of emotional processing outside awareness was observed in the pupillary responses. Previous studies with blindsight patients have shown pupil modulations in response to visual attributes and object categories presented to their blind field (Tamietto et al., 2009; Weiskrantz, Cowey, & Barbur, 1999; Weiskrantz, Cowey, & Le Mare, 1998) that were similar to when stimuli were consciously perceived (Tamietto et al., 2015). For example, Tamietto and colleagues (2015) showed similar pupil dilation increases for consciously perceived and unseen fearful bodies (Tamietto et al., 2015). In line with this, no significant differences were observed between PAS1, corresponding to “no experience”, and the rest of PAS levels within each emotion and time bin. However, we also found no emotion effects during non-conscious body processing (PAS1) in any of the time bins, which was also the case for the rest of PAS levels. It could be that the different analysis procedures of the pupil data may have led to this discrepancy in results with respect to earlier studies.

### Angry and fearful body expressions are processed differently

A central question is how distinct levels of perceptual awareness relate to the different emotional expressions. As mentioned earlier, we observed a gradual increase in sensitivity as the perception of the body expressions became clearer, yet there were some differences between emotions (see **Figure 2A**). While sensitivity values significantly differed between each of the PAS levels for fear, this was not the case between the “no experience” and “brief glimpse” levels of anger. Moreover, the analysis of model slopes indicated that the increase in emotional recognition sensitivity observed with increased subjective awareness was slower for angry body expressions than for fearful ones. These findings indicate that body expression recognition is not only influenced by the general level of perceptual awareness but also by the specific emotion.

Furthermore, higher recognition sensitivity was observed for fearful than angry bodies in all perceptual awareness levels except during perceptual unawareness (i.e., PAS1; see **Figure 2A**). This emotion effect could not be attributed to differences in reaction times between emotions, nor to a bias in the categorization of emotion (**Figure 2B**). A possible interpretation of these findings could be related to the fact that fearful bodies provide more ambiguous information about the locus of the threat than angry bodies, which may have in turn sharpen the discriminability of fearful body expressions over angry ones. In other words, the fact that we observed a higher sensitivity for fearful body expressions suggests a particularly adaptive mechanism essential for disambiguating the self-relevance of a threatening signal and therefore requiring further neural and attentional resources. Similar findings have been reported in studies investigating the effects of gaze on the discrimination of facial expressions. Gaze effects are the strongest for weak, and therefore more ambiguous, facial expressions (Cristinzio, N’diaye, Seeck, Vuilleumier, & Sander, 2010; N’diaye, Sander, & Vuilleumier, 2009) in comparison to faces expressing emotion in a more intense and clear manner (El Zein, Gamond, Conty, & Grèzes, 2015; Graham & LaBar, 2012). Interestingly, gaze direction only modulated this effect once threatening faces were consciously perceived but not outside awareness (Caruana, Inkley, Zein, & Seymour, 2019).

When looking at suppression times, we found that anger body expressions escaped suppression faster than neutral or fearful ones (**Figure 2E**), corroborating the findings by Zhan and colleagues (2015) (Zhan et al., 2015). Here, we extend these findings by showing a consistent effect across different levels of perceptual awareness. As previously mentioned, angry bodies are a much more direct and unambiguous threatening signal, quickly triggering motor preparation responses (i.e., flight/fight behavior). Fearful bodies, on the contrary, convey the nature and source of the threat in a more ambiguous way, prompting the opposite effect in the observer (i.e., freezing responses) (Roelofs, 2017). In line with this, increased motor cortex excitability has been observed for angry bodies compared to fearful or neutral ones (Hortensius, de Gelder, & Schutter, 2016). Neuroimaging experiments have also shown increased activity in the action preparation network in response to angry bodies (Pichon, de Gelder, & Grezes, 2008; Pichon et al., 2009). Taken together, differences in the ambiguity of the threat may thus explain both the sensitivity and breaking from suppression findings: fearful bodies need more attentional resources to disambiguate the threat (expressed as increased sensitivity) while angry bodies may not require such processes and thus trigger faster reactions (expressed as faster breaking from suppression).

With regards to pupil dilation, we found that the influence of body expression on pupil size depends on the breaking from suppression time. Fearful bodies elicited a more constricted pupil size than neutral bodies, but only marginally and for early breaking from suppression times (**Figure 4B**). These findings are in disagreement with previous studies showing larger pupil dilations for negative valenced stimuli than neutral ones (Bradley, Miccoli, Escrig, & Lang, 2008; Kashihara, Okanoya, & Kawai, 2014; Partala & Surakka, 2003). A possible explanation may be that previous studies investigated pupil size modulations by averaging over the whole duration of the trial while in Experiment 2 pupil size changes were only investigated at the moment of breaking from suppression. It may be that pupil dilation has a different time course before and after the body stimulus enters conscious awareness. In fact, in Experiment 1, an increasingly bigger pupil dilation was observed for angry bodies with respect to neutral and fearful ones over time, although not significantly (**Figure 3**).

In conclusion, our results show that behavioral as well as pupillary responses have a gradual relationship with perceptual awareness, but also that this relationship rather than being absolute, was influenced by the specific stimulus category.

## Acknowledgments

This work was supported by the European Research Council (ERC) FP7-IDEAS-ERC (Grant agreement number 295673; Emobodies), by the ERC Synergy grant (Grant agreement 856495; Relevance), by the Future and Emerging Technologies (FET) Proactive Program H2020-EU.1.2.2 (Grant agreement 824160; EnTimeMent) and by the Industrial Leadership Program H2020-EU.1.2.2 (Grant agreement 825079; MindSpaces).

## Author contributions

Conceptualization: M.P.S., M.Z. and B.d.G.; Methodology and Software: M.P.S. and M.Z.; Investigation & Formal Analysis: M.P.S.; Writing – original draft: M.P.S.; Writing – review & editing: M.P.S., M.Z. and B.d.G; Visualization: M.P.S.; Funding Acquisition: B.d.G.

