## Supplementary material for "Gradual relation between perceptual awareness, recognition and pupillary responses to social threat": Table S

† Now at Cognitive Neuroimaging Unit, CEA DRF/I2BM, INSERM, Université Paris-Sud, Université Paris-Saclay, NeuroSpin Center, France

**Table S1. Average estimates of Bayesian Information Criterion values resulting from the gradual and dichotomous model fitting.**

| | Gradual model | Dichotomous model |
|---|---|---|
| **Anger** | -0.52 + 1.14 | 4.19 + 1.15 |
| **Fear** | 0.87 + 1.12 | 6.86 + 1.13 |

Note: the reported values are estimated marginal means and standard errors.

**Table S2. Means and Standard Errors for the reaction times of the Emotion categorization and Perceptual Awareness Scale ratings for each session of Experiment 1.**

| **Emotion categorization** | **HIT** | **MISS** | **FA** | **CR** |
|---|---|---|---|---|
| FN session | 0.84 ± 0.03 | 0.89 ± 0.04 | 0.90 ± 0.04 | 0.88 ± 0.03 |
| AN session | 0.86 ± 0.03 | 0.90 ± 0.03 | 0.91 ± 0.03 | 0.88 ± 0.03 |
| **PAS response** | **PAS1** | **PAS2** | **PAS3** | **PAS4** |
| FN session | 0.58 ± 0.03 | 0.66 ± 0.03 | 0.70 ± 0.03 | 0.65 ± 0.04 |
| AN session | 0.57 ± 0.03 | 0.65 ± 0.03 | 0.72 ± 0.03 | 0.68 ± 0.04 |

PAS: perceptual awareness scale: PAS1: "no experience", PAS2: "brief glimpse"; PAS3: "almost clear experience", PAS4: "clear experience". Hit: emotional stimuli correctly categorized; Miss: emotional stimuli categorized as neutral; False alarms (FA): neutral stimuli categorized as emotional; Correct rejection (CR): neutral stimuli categorized as neutral; FN: fear/neutral; AN: anger/neutral. Note: the reported values (in seconds) are estimated marginal means.

**Table S3. Means and Standard Errors of the reaction times of the Perceptual Awareness Scale ratings of Experiment 2.**

| | PAS1 | PAS2 | PAS3 | PAS4 |
|---|---|---|---|---|
| **Neutral** | 0.85 + 0.08 | 0.74 + 0.08 | 0.78 + 0.08 | 0.80 + 0.08 |
| **Fear** | 0.90 + 0.08 | 0.78 + 0.08 | 0.82 + 0.08 | 0.74 + 0.08 |
| **Anger** | 0.81 + 0.08 | 0.69 + 0.08 | 0.74 + 0.08 | 0.73 + 0.08 |

PAS: perceptual awareness scale: PAS1: “no experience”, PAS2: “brief glimpse”; PAS3: “almost clear experience”; PAS4: “clear experience”. Note: the reported values (in seconds) are estimated marginal means.

**Table S4. Means and Standard Errors of the pupil size during the 2-s CFS period of Experiment 1, separately by Perceptual Awareness Scale rating, emotion and Time bin.**

| | | Neutral | Fear | Anger |
|---|---|---|---|---|
| **PAS1** | **TimeBin1** | -0.01 + 0.01 | 0.00 + 0.01 | 0.00 + 0.01 |
| | **TimeBin2** | -0.32 + 0.03 | -0.35 + 0.06 | -0.31 + 0.05 |
| | **TimeBin3** | -0.30 + 0.04 | -0.32 + 0.07 | -0.31 + 0.06 |
| | **TimeBin4** | -0.21 + 0.05 | -0.25 + 0.08 | -0.21 + 0.06 |
| **PAS2** | **TimeBin1** | 0.00 + 0.01 | 0.00 + 0.02 | -0.01 + 0.02 |
| | **TimeBin2** | -0.30 + 0.03 | -0.32 + 0.03 | -0.30 + 0.03 |
| | **TimeBin3** | -0.28 + 0.05 | -0.29 + 0.05 | -0.27 + 0.05 |
| | **TimeBin4** | -0.17 + 0.05 | -0.17 + 0.06 | -0.20 + 0.05 |
| **PAS3** | **TimeBin1** | 0.02 + 0.02 | 0.00 + 0.01 | 0.02 + 0.02 |
| | **TimeBin2** | -0.24 + 0.04 | -0.31 + 0.04 | -0.23 + 0.05 |
| | **TimeBin3** | -0.19 + 0.05 | -0.26 + 0.05 | -0.2 + 0.07 |
| | **TimeBin4** | -0.09 + 0.06 | -0.12 + 0.06 | -0.08 + 0.08 |
| **PAS4** | **TimeBin1** | 0.00 + 0.02 | -0.05 + 0.01 | -0.01 + 0.02 |
| | **TimeBin2** | -0.29 + 0.03 | -0.24 + 0.04 | -0.29 + 0.06 |
| | **TimeBin3** | -0.26 + 0.05 | -0.20 + 0.05 | -0.23 + 0.07 |
| | **TimeBin4** | -0.18 + 0.06 | -0.11 + 0.06 | -0.15 + 0.06 |

PAS: perceptual awareness scale: PAS1: “no experience”, PAS2: “brief glimpse”; PAS3: “almost clear experience”; PAS4: “clear experience”. Note: the reported values (in mm) are estimated marginal means.
